# A pilot Study: Auditory Steady-State Responses (ASSR) can be measured in human Fetuses using fetal Magnetoencephalography (fMEG)

**DOI:** 10.1101/610303

**Authors:** Dorothea Niepel, Bhargavi Krishna, Eric R. Siegel, Rossitza Draganova, Hubert Preissl, Rathinaswamy B. Govindan, Hari Eswaran

## Abstract

**Background:** Auditory steady-state responses (ASSRs) are ongoing evoked brain responses to continuous auditory stimuli that play a role for auditory processing of complex sounds and speech perception. Transient auditory event-related responses (AERRs) have previously been recorded using fetal magnetoencephalography (fMEG) but involve different neurological pathways. Previous studies in children and adults demonstrated that the cortical components of the ASSR are significantly affected by state of consciousness and by maturational changes in neonates and young infants. To our knowledge, this is the first study to investigate ASSRs in human fetuses.

**Methods:** 47 fMEG sessions were conducted with 24 healthy pregnant women in three gestational age groups (30–32 weeks, 33–35 weeks and 36–39 weeks). The stimulation consisted of amplitude-modulated (AM) tones with a duration of one second, a carrier frequency (CF) of 500 Hz and a modulation frequency (MF) of 27 Hz or 42 Hz. Both tones were presented in a random order with equal probability adding up to 80–100 repetitions per tone. The ASSR across trials was quantified by assessing phase synchrony in the cortical signals at the stimulation frequency.

**Results and Conclusion:** Ten out of 47 recordings were excluded due to technical problems or maternal movements. Analysis of the included 37 fetal recordings revealed a statistically significant response for the phase coherence between trials for the MF of 27 Hz but not for 42 Hz. An exploratory subgroup analysis moreover suggested an advantage in detectability for fetal behavioral state 2F (active asleep) compared to 1F (quiet asleep) detected using fetal heart rate. In conclusion, with the present study it was possible to detect human fetal ASSRs for the first time. These findings warrant further investigations of the developing fetal brain.

## Introduction

The auditory cortex undergoes significant maturational changes during the third trimester (1). Studies of the auditory system are therefore also suited to examine basic principles of fetal brain maturation in general.

Currently, two main methods for direct assessment of fetal brain function are established: fetal functional MRI (2) and fetal MEG (fMEG) (3). Functional MRI measures cerebral blood flow associated with brain activity, and can be mapped with very good spatial, but not temporal, resolution. In terms of the fetal auditory system, a recent functional MRI study in fetuses at the beginning of the third trimester detected a response to auditory stimuli in the primary auditory cortex, indicating sound processing beyond the reflexive, subcortical level (2).

fMEG is completely noninvasive, silent, and has high temporal resolution. We and others have shown successful measurement of transient auditory event related responses (AERRs) (4,5). The detection rate of AERRs is approximately 50% (4,5) but could be increased to 80% in serial measurements (6). Fetal magnetocardiogram (fMCG) is a major by-product of fMEG data (7) that can be used to define fetal behavioral states (8). Previous studies on AERRs to pure tones have shown that AERR latencies decrease with increasing gestational age (9) and increase in fetuses with clinical risk factors such as growth restriction (10) or maternal insulin intolerance (11). In addition, AERRs have been used to further understand basic principles of fetal learning, such as sound discrimination (12), habituation (13) and numerosity discrimination (14). AERRs were also recorded to white noise (15) and to simple syllables (‘ba’ and ‘bi’) (16).

ASSRs are another kind of auditory evoked brain responses that play an important role in auditory processing of complex sounds and speech perception. To our knowledge, ASSRs had never been recorded in the human fetus. The major clinical application are objective electrophysiological hearing tests in adults and young infants (17,18) and monitoring the level of consciousness in general anesthesia (19). ASSRs are evoked by repetitive auditory stimuli or continuous tones with an amplitude and/or frequency modulation at a specific modulation frequency (MF) (20). The cortical response is a continuous waveform at the MF that remains constant in amplitude and phase of its constituent frequency (Fourier) components over a long period of time (21).

AM tones are interesting because of their similarity to phonetic elements that are important for speech perception. Adult MEG studies have shown that stimulation with AM tones can simultaneously elicit two auditory event-related responses: AERRs and ASSRs (22,23). While the present study focuses on fetal ASSRs, a complementary study at the fMEG center in Tübingen (24) investigated AERRs to the same AM tones also used in this study. It was shown that the latencies of AERRs to the onset of AM tones decrease with increasing MF indicating, that the fetus can differentiate between different MFs (24).

The interaction between AERRs and ASSRs is complex. Early studies suggest that ASSRs at 40 Hz are a result of the superimposition of the middle latency components in AERRs (25).

However, not all phenomena observed for ASSRs can be explained with this theory (20) and adult MEG studies revealed that ASSRs and AERRs are localized to different parts of the auditory cortex (22,23). The intrinsic oscillation hypothesis suggests that the ASSR at 40 Hz is generated by oscillation in thalamocortical networks that can synchronize higher auditory associative areas and non-auditory cortices (20,26). The distinction between AERRs and ASSRs is important because it implies different functional applications (26). In the setting of this study, new aspects of the maturation of the fetal auditory system, sound processing and language learning could be investigated. On the other hand, the fact that AERRs have already been recorded using fMEG indicates that the detection of auditory evoked potentials in general is feasible using fMEG.

In awake and relaxed adults, the amplitude of the ASSR decreases with increasing MF in a non-linear pattern (20,27). It is considerably larger for MFs below approximately 50 Hz compared to the higher frequencies. Furthermore, an enhancement of the ASSR can be seen in the range around 40 Hz, and a second (2–5 fold smaller) enhancement occurs between 80 Hz–100 Hz (20,28). The EEG background activity decreases with increasing frequency (20,27,29). Taken together, these facts explain why the most favorable signal-to-noise ratio can be achieved using a MF of 40 Hz–50 Hz, followed by the range between 80 Hz–100 Hz (20,27,29).

It is widely accepted that the ASSR in the region around 40 Hz contains mostly activity from the auditory cortex with only a small proportion of signal coming from the brain stem, whereas the smaller ASSR around 80 Hz predominantly derives from the brain stem (20,26). A common hypothesis is that the higher components of the central auditory nervous system may have less ability to sustain a rhythmic response at higher frequencies (20,26). In line with this, a study using EEG in adults has been shown to localize the ASSR at 88 Hz predominantly to the brain stem with minor subsequent cortical activity. When using a MF of 39 Hz, brainstem activation remained, but a significant cortical activation also occurred (30).

The state of sleep or wakefulness has profound effects on human brain activity, particularly for higher cortical functions. The sleeping state attenuates the ASSR at 40 Hz (predominantly containing signal from the auditory cortex) by approximately 50% (19,20,29). Moreover, the ASSR at 40 Hz decreased in correlation with delta and theta waves, which occur in deeper stages of sleep (32). In contrast, the effect of sleep was much less pronounced in ASSRs to a MF above 70 Hz (predominantly containing signal from the brain stem) (20,29). An advantage of studies of sleeping states is that the EEG background noise is generally lower (20,29). The resulting most efficient MFs varied according to study design and carrier frequency (CF) (33). For CFs of 1000 Hz or below, MFs in the range of 40 Hz or in the range of 90 Hz were reported to be most efficient (29).

Maturational effects also play role for ASSR testing (20,34,35). In neonates, ASSR studies are usually conducted in sedated or unsedated sleep for practical reasons (33,34) so that the results are most comparable to adult studies also conducted during sleep. The amplitude of the ASSR in young infants decreases monotonically with increasing MF (20,36,37). In contrast to adult data (20,29), an enhancement of the ASSR in the range of 40 Hz has not been described (38–40).

Instead, the response at 40 Hz is only half the response at 10 Hz (20) and can not reliably be detected in newborns (41). It was hypothesized that the neonatal auditory cortex was still too immature to sustain a rhythmic response at higher rates (20,34) and that cortical responses may be present at even lower frequencies (38). Previous studies using different methods to detect an ASSRs in neonates and young infants propose an advantage for the higher MFs between 60 Hz–100 Hz that predominantly contain signal from the brain stem (40,54).

The main goal of the current study was to investigate if fetal ASSRs can be measured using fMEG. A thorough literature research did not reveal any previous publications of a successfully recorded ASSR in the human fetus using any technique. We aimed to develop a novel data analysis method taking into account the specific characteristics of the ASSR and the challenges of the fMEG setting. Another focus was to compare the exemplary MFs of 27 Hz and 42 Hz in their suitability for detection of the fetal ASSR. Based on previous fMEG studies using AERRs and previous ASSR studies in neonates and adults, we also hypothesized that maturational effects and effects based on fetal behavioral states may be detectable.

## Materials and Methods

### Participants

We recruited 24 healthy women with an uncomplicated singleton pregnancy. Three groups for gestational age were defined, and each woman could participate once per age group.

- **Group 1:** 30 weeks and zero days until 32 week and six days
- **Group 2:** 33 weeks and zero days until 35 week and six days
- **Group 3:** 36 weeks and zero days until 38 week and six days

Seven women took part in all three fetal sessions, nine women in two fetal sessions, and eight women only participated in one fetal session. This resulted in a total number of 47 fetal sessions. The study was approved by the local institutional review board. Written consent was obtained from every subject.

### Data Acquisition

All data of this study were recorded using a designated fMEG device called the SARA system at the department of Obstetrics and Gynecology at the University of Arkansas for Medical Sciences (SARA = SQUID Array for Reproductive Assessment; CTF Systems Inc., Port Coquitlam, Canada). The surface of the sensor array is shaped in a concave manner to fit the gravid abdomen, and contains 151 primary magnetic SQUID sensors spaced approximately 3 cm apart with a noise level below 5fT/Hz. The sampling rate is 312.5 Hz (5,6). The SARA system is installed in a magnetically shielded room (MSR) (Vakuumschmelze, Hanau, Germany). For the fetal recordings, the pregnant woman was seated in front of the SARA device and advised to lean forward into the sensor array (9). Just prior to every session, a quick sonography was performed to determine the exact position of the fetal head. A fiducial system consisting of four coils was used to mark the position of the fetal head in relation to the SQUID sensors. Before each recording, these coils were activated in a specific frequency and the position of the head coil was calculated (5). All women were instructed to be as still as possible during the recordings, but if a woman became uncomfortable on SARA, the measurements could always be stopped.

### Stimulation

The auditory stimulation of the study was computed using the software Presentation (www.neuobs.com). To avoid magnetic signals correlated to the stimulation, the auditory stimulation was created by speakers outside of the MSR. The tones were then transmitted inside the MSR using an air-filled plastic tubing with an inflated plastic balloon attached to the distal end. During the recording, this balloon was placed between the upper part of the sensor array and the upper abdomen of the participating woman.

The recordings used in this study had a duration of ten minutes, and were conducted as the first recordings of an fMEG session with maximum duration of 30 minutes. Our stimulation paradigm uses AM tones with a modulation depth of 100%, a CF of 500 Hz and a MF of either 27 Hz or 42 Hz. MFs with a known high environmental noise level were avoided (e.g. the power line frequency/first subharmonic in the US at 60 Hz/30 Hz, in the EU at 50 Hz/25 Hz). The (relatively low) CF at 500 Hz has the advantage that it reportedly evokes a larger ASSR at 40 Hz than higher CFs do (42). In addition, lower CFs are attenuated less by the maternal abdomen (43). The volume of the stimulation was set to 94 dB at the surface of the maternal abdomen. Taking into account an estimated attenuation of 20–30 dB (43,44), the volume reaching the fetus is expected to be between 64–74 dB.

The duration of each tone was set to be 1.0 second while the inter-stimulus interval varied randomly between 3.0 seconds and 3.5 seconds, resulting in a stimulus-onset asynchrony of 4.0–4.5 seconds. A randomized process selected a MF of either 27 Hz or 42 Hz with a 50 percent probability prior to every stimulation. This resulted in approximately 80–100 repetitions of each stimulus during a ten-minute recording. Fig 1 visualizes the algorithm as described.

**Fig 1.**
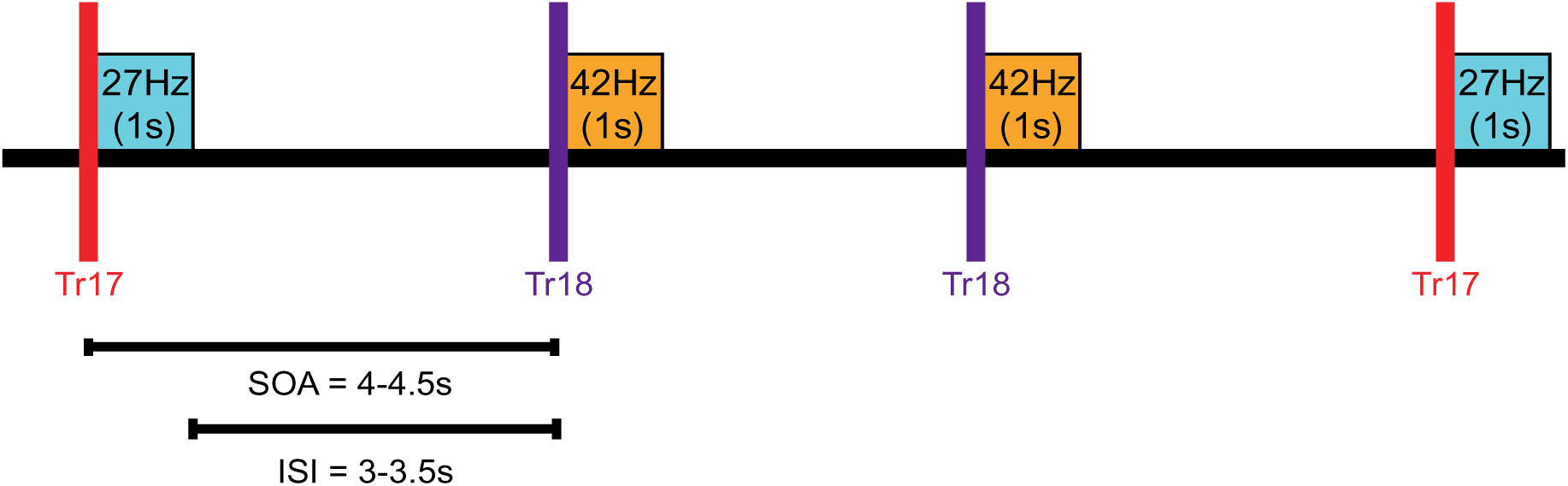
Stimulation paradigm. Illustration of the stimulation paradigm using AM tones with a MF of 27 Hz or 42 Hz in a random sequence. The specific frequency was encoded by a unique trigger value (Tr17 or Tr18). The inter-stimulus interval (ISI) and the stimulus-onset asynchrony (SOA) are displayed.

This protocol is designed to simultaneously evoke AERRs and ASSRs as previously described for adults (22,23). It is very similar to the protocol a complementary study at the fMEG center in Tübingen that investigated AERR to the onset of AM tones and was conducted during a similar time span (24).

### Data Analysis

The preprocessing of the data was done using CTF DataEditor (version: 5.4.0, VSM Med Tech, Coquitlam, BC, Canada). We bandpass filtered the data between 1–50 Hz using the eighth order Butterworth filter with zero-phase distortion. A first-gradient noise reduction was added. The maternal and fetal cardiograms were removed using orthogonal projection (45). Then, the data were divided into one second epochs just prior to the AM tone to define a baseline (‘pre-trigger’) and one second epochs during presentation of the AM tone (‘post-trigger’). A post-trigger paired with its corresponding pre-trigger was defined as trial.

The further data processing was done using MATLAB (version 8.0.0.783(R2012b), The MathWorks, Inc., Natick, MA, USA). Our data analysis utilizes the phase coherence aspect of the ASSR: If the ASSR stimulation elicited a response, the phase of the ASSR at the MF will be synchronous or coherent among all post-trigger segments. The values of phase coherence were calculated for each sensor as follows: The post-trigger and the pre-trigger data were considered for every trial and every sensor and their means were subtracted. A Fourier transform was calculated for MEG in each sensor for each trial and the phase at the investigated MF was calculated using the arctangent transform of the Fourier coefficients. To this, the synchronization index was calculated with a first mode of Fourier transform of the phase. The synchronization index of pre-trigger data was then subtracted from that of the post-trigger data. These phase coherence values were color-coded and plotted as contour maps of the SARA sensor array coordinates. The location of the head coil and the center of activity, a virtual integration of coherence values and their location on the sensor array, were also displayed. The amplitude values at the investigated MF for pre-trigger, post-trigger and post-trigger minus pre-trigger difference were also plotted on contour maps. Negative controls were created within the stimulation recordings by interchanging the markers of the two MFs. That way, the ASSR at 27 Hz was measured during 42 Hz stimulation and vice versa.

For the analysis of fetal behavioral states, the maternal heartbeat was removed using orthogonal projection (45) or a Hilbert Transform algorithm (46). First, the fetal actogram, that displays fetal gross movement, was calculated by tracking the changes in the R-wave amplitude measured from all of the SARA sensors. Then, the cardiogram, that visualizes the fetal heart rate over time, was plotted in a CTG-like fashion. The combined fetal actocardiogram was used to determine the fetal behavioral states by visual inspection. The criteria was based on the Nijhuis criteria (47) and modified according to previous fMEG studies (48,49) as summarized in Table S1. In fetuses 32 weeks or older, behavioral states were be divided in four groups: 1F (passive sleep), 2F (active sleep), 3F (quiet awake) and 4F (active awake). In fetuses with a gestational age below 32 weeks, behavioral states could only be distinguished between ‘passive state’ and ‘active state’. A fetal behavioral state could be identified only if an observed pattern lasted for a minimum of three minutes. This method was well-established at the fMEG center in Tübingen at the time of our study (8,50–52) and all of our results were verified by an expert from the fMEG center in Tübingen.

Our statistical analysis was performed using SAS (version 9.4., The SAS Institute, Cary, NC). For each recording, the pre-trigger and post-trigger values of the phase coherence were considered for all sensors within a 10 cm radius around the head coil. Data were analyzed using a mixed-models repeated-measures approach. In the mixed model, the “subject” was the study participant during a particular session, the within-subjects fixed effect was ‘type of stimulus’ (stimulation or negative control) delivered during the session, and the random effect was the participant considered over all of her sessions. In the group analysis, 90% confidence intervals were calculated for the post-trigger minus pre-trigger values. A p-value below 0.1 was considered statistically significant. In the subgroup analyses, a second fixed effect was added for ‘gestational age group’ or ‘fetal behavioral state’, respectively.

## Results

### Number of Recordings

A total number of 47 recordings was conducted within this study. Ten out of the 47 recordings were excluded: three due to maternal discomfort or extensive movements and seven due to technical problems (wrong or missing head-coil information, wrong or missing stimulation or problems in saving the recording). Table 1 provides an overview of the distribution of recordings between gestational age groups.

**Table 1.** Distribution between gestational age groups.

|  | **Age Group 1**<br>(30w0d-32w6d) | **Age Group 2**<br>(33w0d-35w6d) | **Age Group 3**<br>(36w0d-38w6d) | **All Recordings**<br>(30w0d-38w6d) |
| --- | --- | --- | --- | --- |
| **Total** | 15 | 16 | 16 | 47 |
| (included/excluded) | 11/4 | 14/2 | 12/4 | 37/10 |

**Table 1**. All recordings (total, included, excluded) of this study separated by age group (w = week, d = days).

### Group Analysis

Fig 2 visualizes the results of the overall group analysis for the post-trigger minus pre-trigger phase-coherence values using a MF of 27 Hz or 42 Hz. The exact statistical values are provided in Tables S2 and S3.

**Fig 2.**
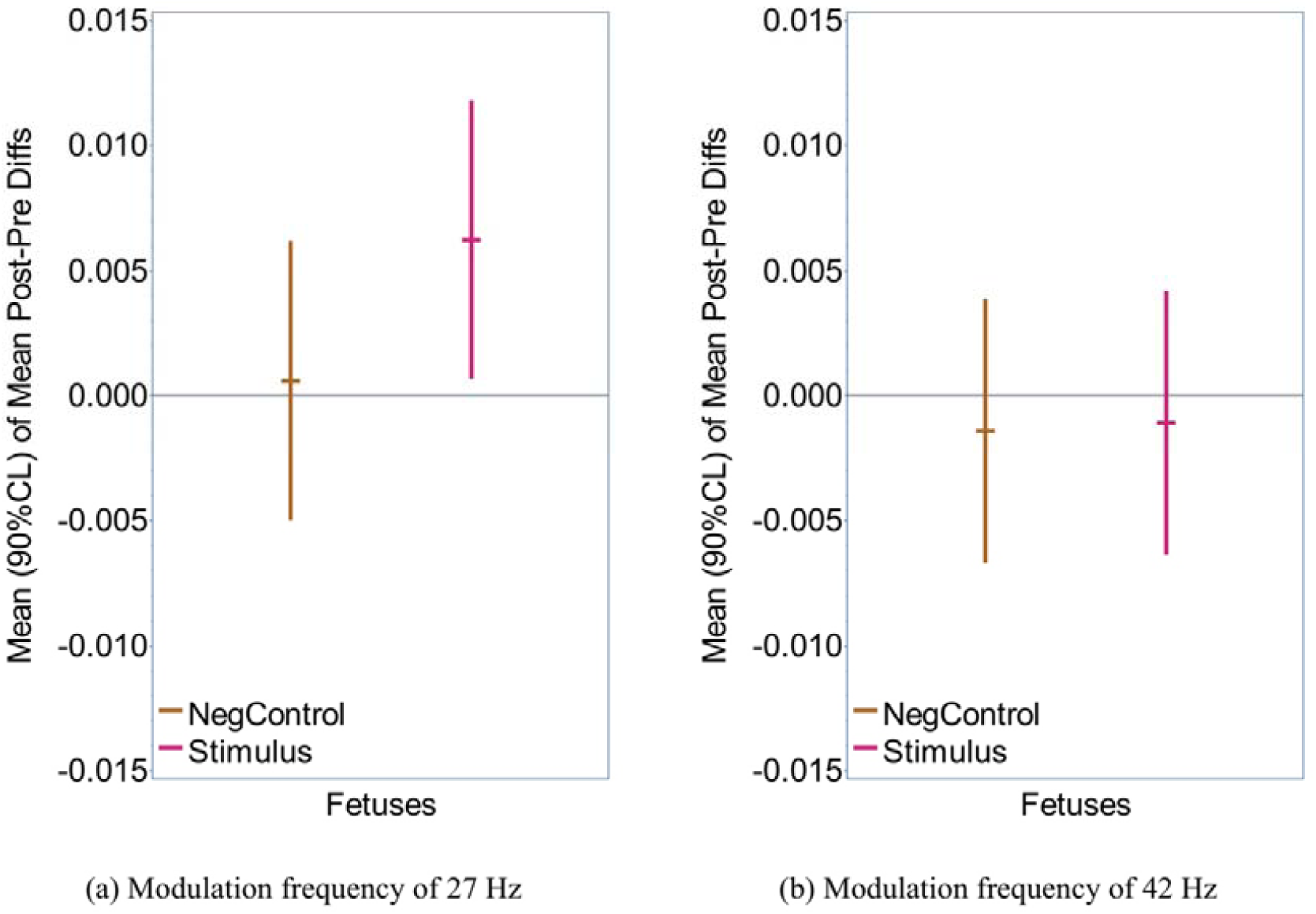
Overall group analysis. Summary of the statistical group analysis of all recordings using a MF of a) 27 Hz, b) 42 Hz. The mean of the post-trigger minus pre-trigger differences of phase coherence is considered for the sensors within a 10-cm radius to the head coil. The error bars represent 90% confidence intervals. For each MF, stimulation recordings (‘Stimulus’) and negative-control recordings (‘NegControl’) are displayed.

A positive results of the post-trigger minus pre-trigger difference means that the level of phase coherence is higher in the post-trigger data (during stimulation) than in the pre-trigger data (prior to stimulation). In theory, a negative result means that the pre-trigger data was more coherent at the MF.

The most important finding in this study is that the analysis of the 27 Hz ASSRs has a significantly positive result (p-value = 0.0667) for the stimulation recordings (n = 37, pink line in figure 2a). These findings are further confirmed by the fact that the negative controls (n = 37; brown line in figure 2a) were not significantly different from zero. In contrast, the overall analysis of the ASSR to the MF of 42 Hz did not reveal any statistically significant results (n = 37; figure 2b).

### Individual Responses

Within the stimulation recordings using the MF of 27 Hz, nine out of 37 (24.3%) recordings had a statistically significant positive response, five (13.5%) had a negative result and 23 (62.2%) were not significantly different from zero. In the negative controls, six out of 37 recordings (16.2%) had a positive response, three (8,1%) were negative and 28 (75.7%) were not significantly different from zero.

### Contour Maps

In the next step, we computed contour maps of the SARA sensor array in order to visualize the distribution of the signals investigated. Fig 3 shows the contour maps of two recordings using amplitude and phase-coherence values at a MF of 27 Hz for pre-trigger, post-trigger, and post-trigger minus pre-trigger data.

**Fig 3.**
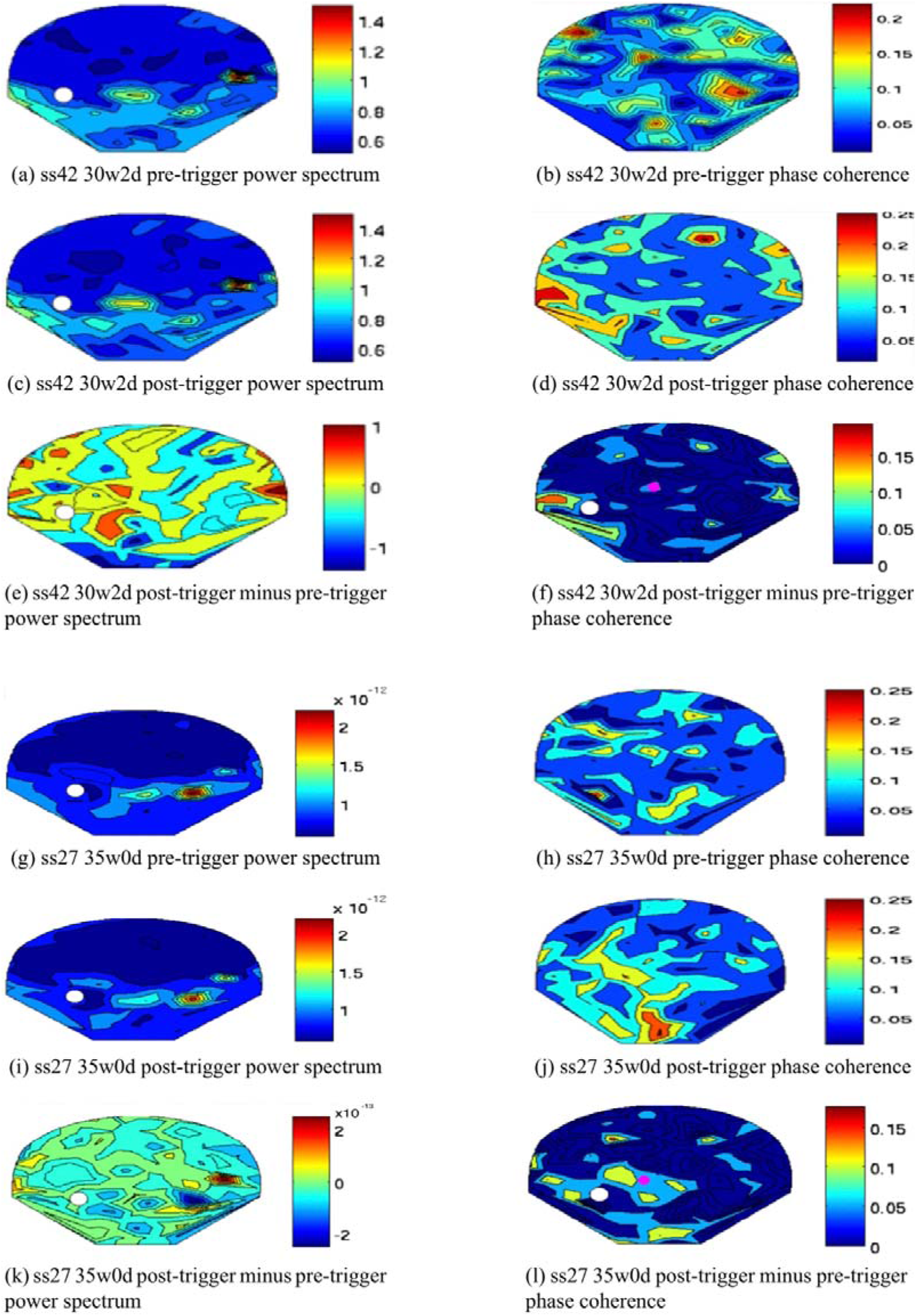
Contour maps. Comparison between the amplitude approach (‘power spectrum’) and the phase coherence approach visualized on contour maps of the SARA sensor array. Subfigure (a)–(f) and subfigure (g)–(l) have been computed from two different recordings (‘ss37’ and ‘ss27’). Gestation age is also displayed (w=weeks, d=days). The white circles represent the head coil, the pink circles in subfigure (f) and (l) represent the center of activity.

In the 27 Hz phase-coherence maps, an activation close to the head coil appeared to be present in a proportion of the post-trigger and the post-trigger minus pre-trigger (see subfigure (d), (f), (j) and (l)). Although the origin of this activation cannot be objectively determined, an ASSR would be expected to cause an increased phase coherence close to the head coil during stimulation (post-trigger) compared to baseline (pre-trigger). Therefore, it appears possible that the observed signal may represent fetal ASSR, although other causes including random events cannot be ruled out.

It is also remarkable that the pre-trigger maps in the amplitude approach each seem to demonstrate a specific pattern that is very similar in the corresponding post-trigger map. These patterns may represent specific physiological background activity. For example, multiple recordings exhibited an activation in the lower left and lower right sensors similar to the pattern in subfigure (a) and (c). This activation may be due to muscle activity in the maternal legs as a result of the sitting position on SARA. The activity in the center right side of subfigure (a), (c), (g) and (i) could potentially be attributed to fMCG residuals. The phase-coherence maps of the same recordings, on the other hand, show no specific common pattern in the pre-trigger data (see subfigure (b) and (h)). Instead, the baseline activity appears to be randomly distributed throughout the sensor array.

### Effects of Gestational Age

The results of an exploratory subgroup analysis depending on fetal age groups are summarized in Fig 4. The exact statistical values are provided in Table S4 and the sample size of the subgroups can be reviewed in above Table 1.

**Fig 4.**
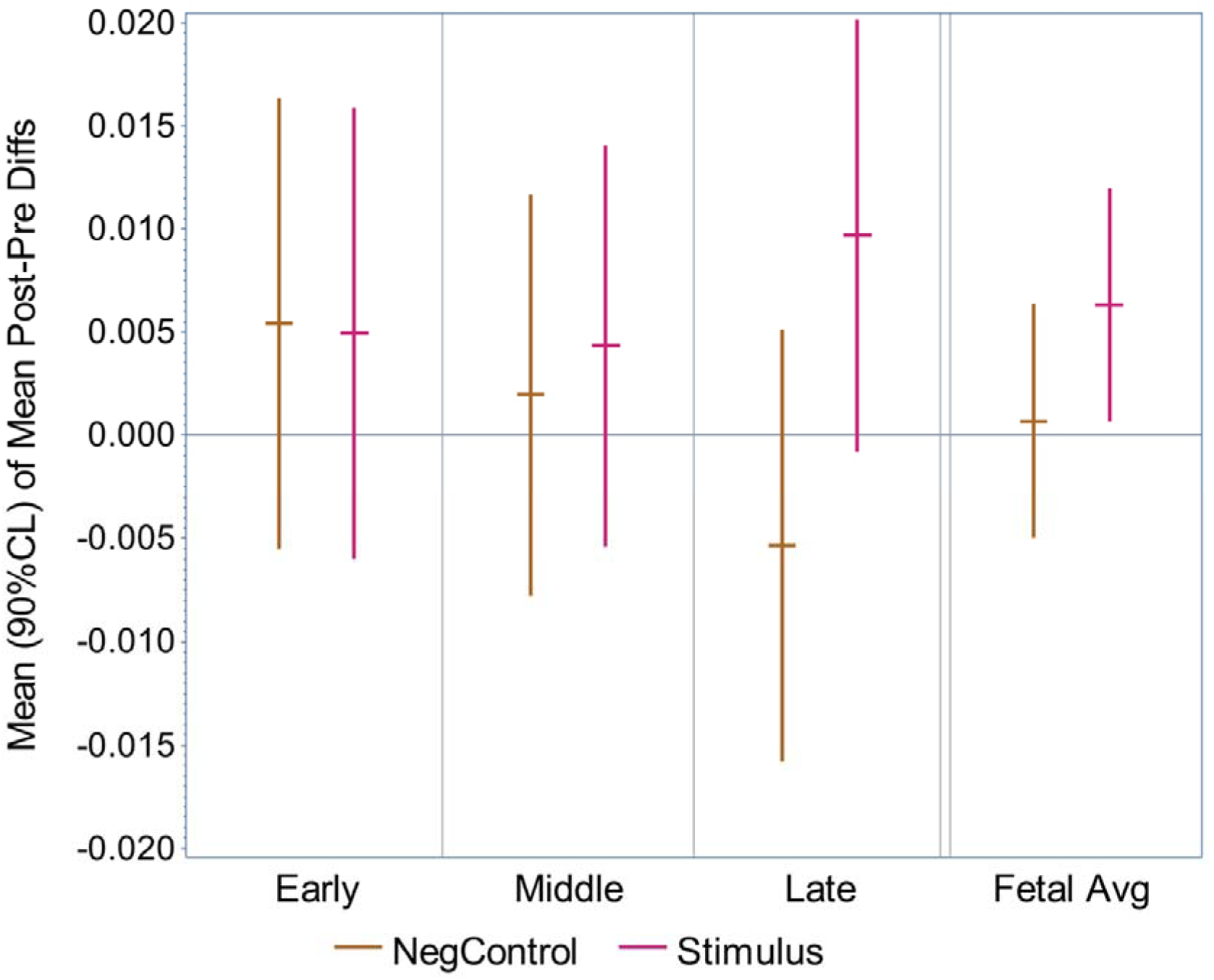
Subgroup analysis of gestational age. Summary of the statistical subgroup analysis depending on gestational age groups using a MF of 27 Hz. The error bars represent the 90% confidence intervals. Gestational age group 1 (‘early’), group 2 (‘middle’) and group 3 (‘late’) and their average (‘Fetal Avg’) are shown. Stimulation recordings (‘Stimulus’) and negative control recordings (‘NegControl’) are displayed.

For the different age groups we did not observe a significant effect between stimulus and negative control condition.

### Effects of fetal Behavioral State

The distribution of the fetal behavioral states among all recordings of this study is demonstrated in Table 2. It is important to consider that the sample size of the various behavioral state subgroups varies significantly (n = 1–15). We thus decided to perform an exploratory analysis only considering the two largest subgroups (2F with n = 15 and 1F with n = 6). All other subgroups had a very small sample size (n = 1–4) and were summarized as “others”. The results are summarized in figure 5 and the exact statistical values are provided in Table S5.

**Table 2.** Distribution between behavioral states.

|  | Age Group 1 | Age Group 2 | Age Group 3 | All Age Groups |
| --- | --- | --- | --- | --- |
| Active State | 4 | n/a | n/a | 4 |
| Passive State | 2 | n/a | n/a | 2 |
| Active State → Passive State | 1 | n/a | n/a | 1 |
| Passive State → Active State | 1 | n/a | n/a | 1 |
| State 1F | 1 | 4 | 1 | 6 |
| State 2F | 2 | 7 | 6 | 15 |
| State 4F | 0 | 0 | 1 | 1 |
| State 1F → 2F | 0 | 3 | 0 | 3 |
| State 2F → 1F | 0 | 0 | 3 | 3 |
| State 3F → 2F | 0 | 0 | 1 | 1 |
| All Recordings | 11 | 14 | 12 | 37 |

**Table 2**. Distribution of the fetal behavioral states among the different gestational age groups. Recordings *<* 32 weeks 1 day of gestation are plotted separately in the top rows because of the different classification system of fetal behavioral states. Behavioral state groups that did not occur within this study are not displayed.

**Fig 5.**
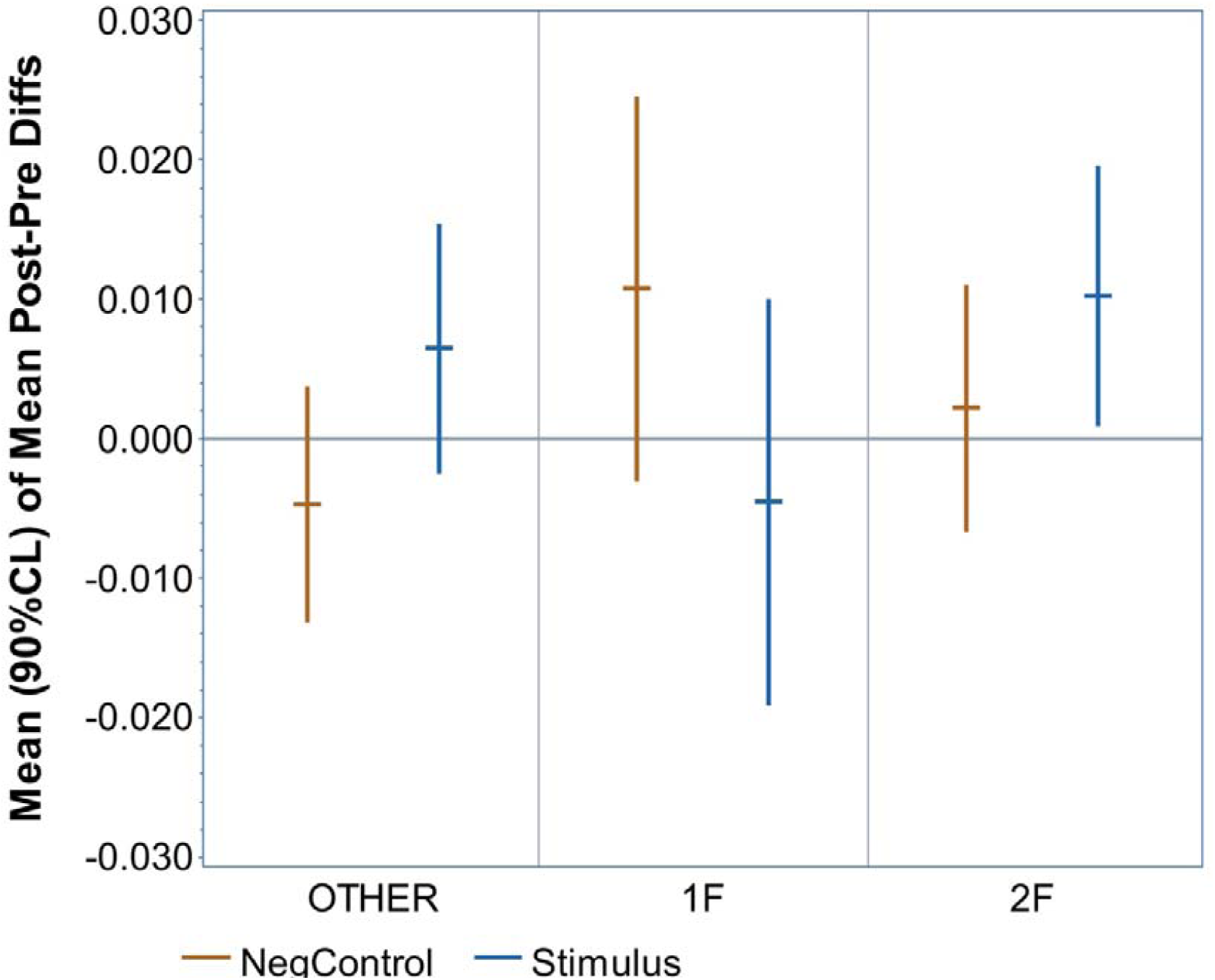
Subgroup analysis of fetal behavioral state. Summary of the statistical subgroup analysis depending on fetal behavioral-state groups using a MF of 27 Hz. The error bars represent the 90% confidence intervals. The most frequent fetal states ‘2F’ (active asleep) and ‘1F’ (quiet asleep) are displayed, all less common fetal states were summarized as ‘OTHER’. Stimulation recordings (‘Stimulus’) and negative control recordings (‘NegControl’) are displayed.

The most important finding is that the stimulation recordings have a significantly positive result (p = 0.0716) in fetal behavioral-state group 2F, the largest subgroup (n = 15). It is also remarkable that the first recordings in group 1F (n = 6) remain close to zero. These findings are further validated by the fact that none of the groups of negative controls were significantly different from zero.

## Discussion

The results of our overall group analysis clearly show a statistically significant response of the stimulation recordings using a MF of 27 Hz while the negative controls were not significantly different from zero (Fig 2). Out of all our statistical analyses, this group has the largest sample size and hence the largest statistical power. Of note, the AM tones used for simulation do not contain a frequency contributions at the MF but only at the CF and at ‘side bins’ at the CF ± the MF (in our case at 500 Hz +/– 27 Hz) (53). The data analysis was moreover completely automated so that the results are truly objective. For all those reasons, it appears likely that the recorded signal reflects a fetal ASSR.

An important focus of the present study was to develop a method for the data analysis of ASSR testing in the fetal setting. Previous adult studies have shown that data analysis using the phase coherence aspect or the amplitude aspect of an ASSR provide similar results in terms of sensitivity and specificity (53). However, it appears plausible that this is different when using fMEG. As demonstrated in the contour maps of the SARA sensor array, relevant physiological background activity at 27 Hz occurs in the amplitude approach but not when using phase coherence (see figure 3). Given that the fetal brain signal is expected to be very small, the amplitude of physiological background activity at 27 Hz may be larger than the investigated ASSR by multiple orders of power. The phase coherence approach on the other hand requires the signal to be correlated to the stimulation in order to be synchronous throughout all trials. Even if environmental signals can change the phase in some trials, the overall effect will be less severe, because the recorded phase in a trial does not reflect the total strength of the signal.

Another main finding of this study is, that a detectable fetal ASSR can be evoked by a MF of 27 Hz, but not by a MF of 42 Hz. A number of physiological factors can in theory explain why the signal of the ASSR is expected to be larger at 27 Hz. A large study in neonates described that although the amplitude of the neonatal ASSR was largest at 25 Hz (the lowest MF investigated) and decreases with increasing MF, this effect was offset by the longer time needed to evoke a response. Parameters such as ‘time to criterion’ and ‘signal strength’ measured by magnitude-squared coherence were more favorable in the middle and high MFs (41 Hz–88 Hz) (36). Remarkably, this advantage was not significant for lowest CF tested at 500 Hz, because the difference in amplitude was more pronounced in the lower CFs (36). This is particularly interesting because the present study is also using a CF of 500 Hz. In the fMEG setting it also has to be considered that the fetal brain signal is much smaller and that the strength of the fetal ASSR may be close to the detection limit. Hence, it appears possible that the higher amplitudes in the lower MFs are necessary to enable detection when using fMEG.

The enhancement of the ASSR around 40 Hz (mostly deriving from the cortex) in adult data has also not been described in studies on neonates or infants (38–40). An early study suggests that, based on the superimposition theory for the generation of ASSRs, the increased latencies of AERRs in young children explain why an superimposition of the peaks is not possible at 40 Hz but can occur, to some extent, for MFs between 20–30 Hz. Accordant with that theory, the authors observed a small increment in the amplitude of the ASSR of young children between 20 Hz and 30 Hz (38).

The fMEG study that used a stimulation protocol similar to the one in this study to investigate AERRs to the onset of AM tones reported that the mean latencies were shortest for the MF of 27 Hz compared to an MF of 2 Hz, 4 Hz, 8 Hz, 42 Hz, 78 Hz and 91 Hz (24). Moreover, and in line with the findings from our study, the detection rates for AERRs were highest at 86% for a MF of 27 Hz compared to 50% for a MF of 42 Hz (24) and approximately 50% in previous studies using pure tones (5). The authors suggest, that the increased detection rates with a MF 27 may be due to a better transmission by bone conduction, the main way of fetal hearing in utero (24).

The distribution of baseline activity should also be considered. In neonatal EEG data, the background activity decreases with increasing MF (36,37). In the fMEG setting, additional physiological and environmental signals are present (e.g. maternal MCG: 1–1.5 Hz, fMCG: 2–3 Hz, muscle contraction: 20–300 Hz (55), power line frequency in the US/EU: 60 Hz/50 Hz) and the resulting background activity varies in a noncontinuous pattern that can be subject to the local setup. For example, a relevant number of recordings in our study exhibit a visible continuous 20 Hz signal possibly connected to noise from a technical device (e.g. the camera frame rate). The first harmonic of this signal at approximately 40 Hz could in theory have distorted our 42 Hz measurements. These examples demonstrate, that each MF has to be evaluated individually in the fMEG setting.

The exploratory subgroup analysis dependent on fetal gestational age showed a tendency towards a positive result in age group 3 (p=0.13) but no statistically significant responses. In theory, this trend supports the hypotheses, that the fetal auditory cortex in the majority of the fetuses in age group one and two may still be too immature to generate an ASSR to a MF of 27 Hz. However, the data are limited because the sample size of the subgroups is relatively small (n = 11–14). Thus, it is not possible to draw a final conclusion regarding potential maturational effects based on this data alone. Investigations of AERRs to the onset of AM tones showed a change in latencies depending on gestational age for the MF of 4 Hz. In line our study, the results were not significant when using a MF of 27 Hz but the sample size of this group was also only half the size of the 4 Hz group (n^(27 Hz)^ = 14 compared to n^(4 Hz)^ = 28) (24). Further studies are needed in order to evaluate potential maturational effects of the fetal ASSR.

The subgroup analysis based on fetal behavioral states seems to suggest a preferable detectability of the ASSR at 27 Hz in fetal state 2F (active asleep) compared to state 1F (quiet asleep). It has to be considered that this data are very limited due to the small sample size of the subgroups. In previous adult studies, the cortical component of the ASSR at and below 40 Hz decreases in the state of sleep (19,25,29) and even more in deeper stages of sleep (32). In line with our observations, an early fetal study that observed changes in heart-rate pattern and fetal movement following auditory stimulation reported more frequent responses in fetal state 2F compared to fetal state 1F (56). A previous fMEG study also demonstrated that the latencies of AERRs is significantly shorter in active sleep (2F) compared to passive sleep (1F) (50). In summary, our data are limited but the fact that our observations are in agreement with previous studies make a correlation between fetal state and fetal ASSR more likely. Future studies should therefore investigate the fetal ASSR in the context of the fetal state.

One of the main challenges of the present study was a low signal-to-noise ratio. The ASSRs could be detected in a group analysis, but not in a significant proportion of the individual recordings. One limitation is that all sensors within a 10 cm radius around the head coil were included in the analysis. In fetal AERR studies, the sensors with the largest amplitude (usually approx. five sensors) are determined by visual inspection and used for further analysis. Although the connection between AERRs and ASSRs is complex and ASSRs may not be produced in the exact same neurological pathways, it is likely that the ASSR is also only present in a few sensors. By including all sensors within 10 cm around the head coil, the ASSR might average out while sometimes, the signal may even be more far away than 10 cm. It would be interesting to select the sensors containing the largest AERR and to use them for an analysis of ASSRs in future studies. A number of more general adjustments to optimize the signal-to-noise ratio in ASSR testing have also been studied in adult data (27,53). It is for example known, that a higher volume of the stimulus increases the ASSR evoked (20,25,27). The present study is using a relatively low volume of 94 dB at the surface of the maternal abdomen. Previous studies using the SARA device have been using a volume of up to 120 dB (5,6). The signal-to-noise ratio could also be improved by increasing the number of trials investigated. This could for example be achieved by increasing the time of the recording, by decreasing the ISI between trials or also by just using one MF per recording instead of two alternating MFs.

## Conclusion

In the present study, it was possible to detect the human fetal ASSR for the first time using fMEG. We demonstrated that a MF of 27 Hz is suitable to evoke a recordable fetal ASSR. With regards to the data analysis, we illustrated that the phase coherence approach has notable advantages over the amplitude approach in the fMEG setting. A limitation of this study is, that the ASSR could only be objectively recorded in a group analysis. A preliminary sub-analysis of our data suggests an advantage for fetal behavioral state 2F (active asleep) compared to fetal behavioral state 1F (quiet asleep) but could not detect significant changes based on gestational age. Future studies are needed to improve the signal-to-noise-ratio and increase the detection rate for individual recordings. It would also be interesting to further investigate the effects of fetal brain maturation and behavioral states to the fetal ASSR. A long-term goal is to establish a method for non-invasive fetal brain surveillance, to detect imminent brain damage early and to put targeted preventive or therapeutic measures into place.

## Supporting Information

**S1 Table: Criteria for the definition of fetal behavioral states.**

This table summarizes the criteria used in this study to define fetal behavioral states based on a visual analysis of the fetal actocardiogram.

**S2 Table: Overall group analysis for the MF of 27 Hz.**

This table provides the exact values of the statistical analysis including the calculated standard error and p-values.

**S3 Table: Overall group analysis for the MF of 42 Hz.**

This table provides the exact values of the statistical analysis including the calculated standard error and p-values.

**S4 Table: Subgroup analysis according to gestational age group for the MF of 27 Hz.**

This table provides the exact values of the statistical analysis including the calculated standard error and p-values.

**S5 Table: Su**b**group analysis according to fetal behavioral state group for the MF of 27 Hz.**

This table provides the exact values of the statistical analysis including the calculated standard error and p-values.

## Acknowledgements

We thank the team of the SARA lab for recruiting and the subjects and maintaining the SARA device. We also gratefully thank all women for their participation.

