## Supplementary material for "A pilot Study: Auditory Steady-State Responses (ASSR) can be measured in human Fetuses using fetal Magnetoencephalography (fMEG)": S1 Table

### S1 Table: Criteria for the definition of fetal behavioral states. This table summarizes the criteria used in this study to define fetal behavioral states based on a visual analysis of the fetal actocardiogram.

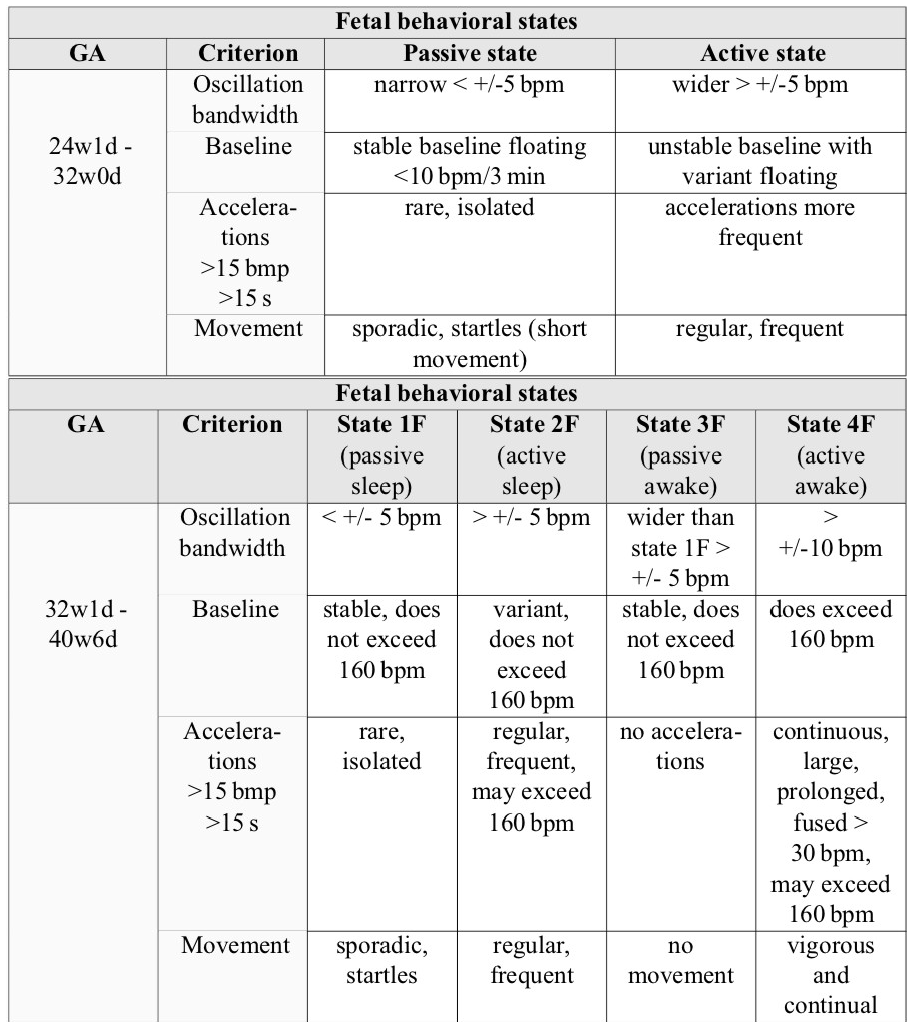

**Table S1.** Criteria used for the definition of fetal behavioral states based on the visual evaluation of the fetal actocardiogram. A state was only defined it a pattern lasted for at least three minutes. bpm stands for beats per minute. Source: MEG Center Tübingen
