## Supplementary material for "A pilot Study: Auditory Steady-State Responses (ASSR) can be measured in human Fetuses using fetal Magnetoencephalography (fMEG)": S3 Table

**S3 Table: Overall group analysis for the MF of 42 Hz.** This table provides the exact values of the statistical analysis including the calculated standard error and p-values.

| **Least Squares Means** | | | | | | | | | |
| --- | --- | --- | --- | --- | --- | --- | --- | --- | --- |
| **Effect** | **reclabel** | **Estimate** | **Standard Error** | **DF** | **t Value** | **Pr > \|t\|** | **Alpha** | **Lower** | **Upper** |
| **reclabel** | NegControl | -0.00139 | 0.003152 | 52 | -0.44 | 0.6602 | 0.1 | -0.00667 | 0.003884 |
| **reclabel** | Stimulus | -0.00106 | 0.003152 | 52 | -0.34 | 0.7377 | 0.1 | -0.00634 | 0.004217 |

**Table S3.** Results of the statistical group analysis for all recordings using the MF of 42 Hz. The estimated post-trigger minus pre-trigger value (‘Estimate’), the corresponding standard error, the degree of freedom (‘DF’), the t-value (‘t Value’), the p-value (‘Pr *> |*t| ’), the significance level (‘Alpha’) and the upper and lower limit of the 90% intervals (‘Upper’ and ‘Lower’) are displayed for stimulation recordings and negative controls.
