## Supplementary material for "A pilot Study: Auditory Steady-State Responses (ASSR) can be measured in human Fetuses using fetal Magnetoencephalography (fMEG)": S4 Table

**S4 Table:** **Subgroup analysis according to gestational age group for the MF of 27 Hz.** This table provides the exact values of the statistical analysis including the calculated standard error and p-values.

| **Estimates** | | | | | | | | |
| --- | --- | --- | --- | --- | --- | --- | --- | --- |
| **Label** | **Estimate** | **Standard Error** | **DF** | **t Value** | **Pr > \|t\|** | **Alpha** | **Lower** | **Upper** |
| **NegControl, Early** | 0.005417 | 0.006524 | 48 | 0.83 | 0.4105 | 0.1 | -0.00553 | 0.01636 |
| **NegControl, Late** | -0.00534 | 0.006252 | 48 | -0.85 | 0.3976 | 0.1 | -0.01582 | 0.005150 |
| **NegControl, Middle** | 0.001974 | 0.005798 | 48 | 0.34 | 0.7349 | 0.1 | -0.00775 | 0.01170 |
| **NegControl, Avg.** | 0.000685 | 0.003382 | 48 | 0.20 | 0.8404 | 0.1 | -0.00499 | 0.006357 |
| **Stimulus, Early** | 0.004940 | 0.006524 | 48 | 0.76 | 0.4526 | 0.1 | -0.00600 | 0.01588 |
| **Stimulus, Late** | 0.009696 | 0.006252 | 48 | 1.55 | 0.1275 | 0.1 | -0.00079 | 0.02018 |
| **Stimulus, Middle** | 0.004332 | 0.005798 | 48 | 0.75 | 0.4586 | 0.1 | -0.00539 | 0.01406 |
| **Stimulus, Avg.** | 0.006323 | 0.003382 | 48 | 1.87 | 0.0677 | 0.1 | 0.000650 | 0.01200 |

**Table S4.** Results of the subgroup analysis according to gestational age groups using the MF of 27 Hz. The estimated post-trigger minus pre-trigger value (‘Estimate’), the corresponding standard error, the degree of freedom (‘DF’), the t-value (‘t Value’), the p-value (‘Pr *> |*t|’), the significance level (‘Alpha’) and the upper and lower limit of the 90% intervals (‘Upper’ and ‘Lower’) are displayed for stimulation recordings and negative controls.
