## Supplementary material for "A pilot Study: Auditory Steady-State Responses (ASSR) can be measured in human Fetuses using fetal Magnetoencephalography (fMEG)": S5 Table

### S5 Table: Subgroup analysis according to behavioral state group for the MF of 27 Hz. This table provides the exact values of the statistical analysis including the calculated standard error and p-values.

**Table S5.** Results of the subgroup analysis according to fetal behavioral state groups of fetuses at or above 32 weeks of gestation using the MF of 27 Hz. The estimated post-trigger minus pre-trigger value (‘Estimated Group Mean’), the corresponding standard error, the degree of freedom (‘DF’), the t-value (‘t-statistic’), the p-value (‘P value’), the significance level (‘Alpha’) and the upper and lower limit of the 90% confidence intervals (‘Lower 90% Conf.Limit on Estimate’ and ‘Upper 90% Conf.Limit on Estimate’) are displayed for stimulation recordings and negative controls.

| **Fetal State** | **reclabel** | **Estimated Group Mean** | **Standard Error of Estimate** | **DF** | **t-statistic** | **P value** | **Alpha significance level** | **Lower 90% Conf.Limit on Estimate** | **Upper 90% Conf.Limit on Estimate** |
| --- | --- | --- | --- | --- | --- | --- | --- | --- | --- |
| 1F | NegControl | 0.01076 | 0.00823 | 48 | 1.307 | 0.1976 | 0.10 | -.00305 | 0.02457 |
| 1F | Stimulus | -.00453 | 0.00870 | 48 | -0.521 | 0.6048 | 0.10 | -.01913 | 0.01006 |
| OTHER | NegControl | -.00474 | 0.00506 | 48 | -0.936 | 0.3538 | 0.10 | -.01322 | 0.00375 |
| OTHER | Stimulus | 0.00646 | 0.00535 | 48 | 1.208 | 0.2331 | 0.10 | -.00251 | 0.01542 |
| 2F | NegControl | 0.00220 | 0.00527 | 48 | 0.417 | 0.6786 | 0.10 | -.00665 | 0.01104 |
| 2F | Stimulus | 0.01025 | 0.00556 | 48 | 1.842 | 0.0716 | 0.10 | 0.00092 | 0.01958 |
